# Hypsochromic shift of phyC complements the inhibition of hypocotyl elongation under moderate red/far-red light

**DOI:** 10.1101/2024.03.08.584055

**Authors:** Shizue Yoshihara, Koji Okajima, Satoru Tokutomi

**Author notes:** **Email address for each author** S. Yoshihara K. Okajima S. Tokutomi.

## Abstract

Phytochrome (phy) is a plant photoreceptor that regulates various photomorphogenesis, and occurs in two forms, a red light (R)-absorbing form (Pr) and a far-red light (FR)-absorbing form (Pfr). Absorption spectral analyses of the photosensory module (PSM) showed that phyC in the Pr of *Arabidopsis thaliana*, *Solanum lycopersicum* and *Zea mays* exhibited the absorption maxima shift toward shorter wavelengths (hypsochromic shift) compared with those of phyA and phyB. Substitution of the chromophore-binding domain complemented the hypsochromic shift in the spectra of phyC in the Pr. The effect of the hypsochromic shift on the inhibition of hypocotyl elongation was studied under R/FR ratio from 0.5-10. PhyB was revealed to play a major role in inhibition, and phyC showed a complementary role under R/FR <2.0. This may result from the activation peak of the phyC PSM, which was hypsochromically shifted compared with that of the phyB PSM from the Pr to Pfr. The leaf-filtered light measurement suggested that phyC enables plants to receive more R and contributes to survival in the field. Under low R/FR conditions, the activation efficiency of phyC was greater than that of phyB, suggesting that the hypsochromic shift of phyC is necessary for the robust growth of angiosperms.

**Highlight:** Angiosperm phyC with hypsochromically shifted activation inhibit hypocotyl elongation under relatively low red/far-red light conditions, in which phyB is not fully functional.

## Introduction

In environments with variable light, plants survive by constantly monitor changes to optimize growth and flowering. For these purposes, plants have acquired and utilized multiple photoreceptors to sense UV-B, blue and red/far-red light (Galvão and Fankhauser, 2015). Phytochrome (phy) is the first photoreceptor found in plants and reversibly converts between an inactive red light (R)-absorbing form (Pr) and an active far-red light (FR)-absorbing form (Pfr); through this conversion, phytochromes regulate a wide variety of plant developmental transitions, such as seed germination, de-etiolation, flowering timing, shade avoidance and integration with thermal and hormonal signals (Legris *et al*. 2019).

The domain architecture of plant phytochromes is highly conserved among higher plants (Rockwell *et al*., 2006) and consists of an N-terminal photosensory module (PSM) with N-terminal extension (NTE) and Per-Arnt-Sim (PAS)-cGMP phosphodiesterase/adenylyl cyclase/FhlA (GAF)-phytochrome (PHY) domains, as well as a C-terminal regulatory module with a PAS-PAS-histidine kinase-related domain (HKRD) (Fig. 1A). A linear tetrapyrrole chromophore, phytochromobilin (PΦB), is covalently attached to a conserved Cys residue in the GAF domain. The PSM plays a major role in reversible photoconversion and signal transduction and has been used to study the photochemistry and physiological functions of phytochromes (Matsushita *et al*., 2003; Su and Lagarias, 2007; Oka *et al*., 2008; Zhang *et al*., 2013; Burgie *et al*., 2014).

**Fig. 1.**
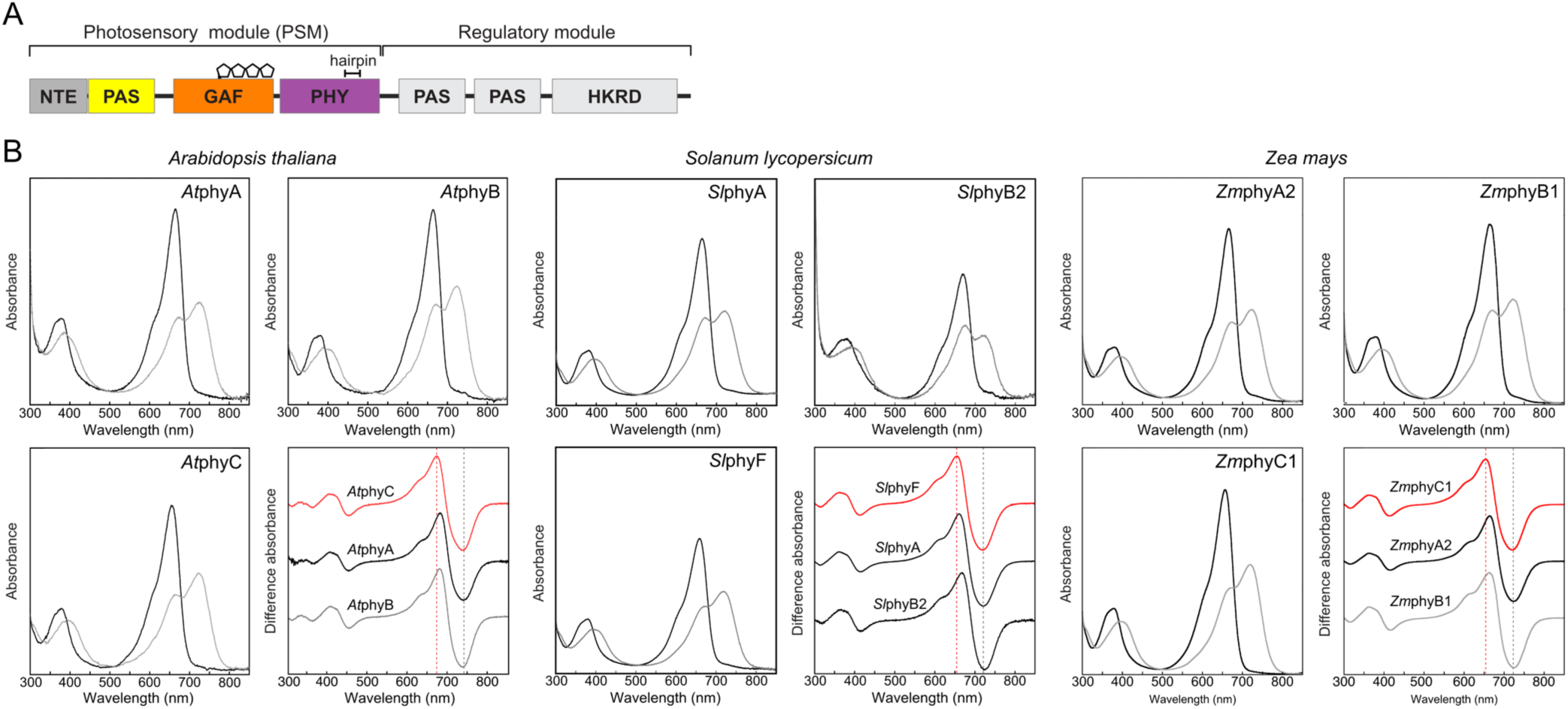
Absorption properties of phytochrome photosensory modules. (A) A typical domain structure of plant phytochrome consisting of an NTE (N-terminal extension) and PAS (Per/ARNT/Sim), GAF (cGMP phosphodiesterase/adenylyl cyclase/FhlA), PHY (phytochrome) domains and HKRD (histidine kinase related domain). The four pentagons of the GAF domain represent the PΦB chromophore. (B) Pr and Pfr absorption spectra and absorption difference spectra between the Pr and the Pfr of photosensory modules (PSMs) of phyA, phyB and phyC-type phytochromes from *Arabidopsis thaliana* (*At*), *Solanum lycopersicum* (*Sl*) and *Zea mays* (*Zm*). The red and gray dashed lines in difference spectra indicate the position of the λ_max_ of phyC-type phytochromes in the Pr and Pfr, respectively.

Molecular phylogenetic analysis suggested that phytochrome genes first diverged into two families containing the ancestral forms of *PHYA* and *PHYC*, as well as *PHYB* and *PHYE* (Mathews, 2006). Subsequent gene duplication yielded the *PHYA* and *PHYC* subfamilies from the former family. *PHYE* may branch from the *PHYB* subfamily in some plant species, and *PHYD* may be duplicated from *PHYB* in *Arabidopsis thaliana* (*At*) (Clack *et al*., 1994).

*PHYA*, *PHYB* and *PHYC* are widely conserved among angiosperms. In *Arabidopsis*, phyA is a dominant phytochrome in etiolated plants and is unstable under light irradiation. Compared with phyA, phyB is a major phytochrome that is active in light-grown plants and is more stable under light conditions (Howe *et al*., 1998). There are many studies on the physiological functions of phytochromes, and among the three phytochromes, phyC is currently the least understood.

In *Arabidopsis*, phyC is reportedly responsible for R sensing. Compared to the WT, seedlings of the *AtphyC* mutant Col-0 background had longer hypocotyls and smaller cotyledons under continuous red light (Rc), in which phyC expression was phyB dependent (Monte *et al*., 2003). Similar phyB-dependent expression of phyC was reported by Hu *et al*. in 2013. Furthermore, phyC senses day length independently of phyB in *Arabidopsis*. Seedlings of *AtphyB* or *AtphyC* mutants flowered early under short-day conditions, but flowering in *AtphyB phyC* double mutant was not further accelerated (Monte et al., 2003). The acceleration of flowering by phyC has also been reported for the monocot plant wheat, although it is under long-day photoperiod (Chen *et al*., 2014).

On the other hand, phyC does not participate in FR sensing in *Arabidopsis*. *AtphyC* mutant seedlings were indistinguishable from WT seedlings under constant FR (FRc) (Monte *et al*., 2003). In contrast, phyC senses FR in the monocot rice. The *phyA* mutant of *Oryza sativa* (*Os*) showed incomplete de-etiolation under FRc, while the *OsphyA phyC* double mutant showed no de-etiolation. Thus, both *Os*phyA and *Os*phyC are involved in FR sensing in rice (Takano *et al*., 2005). The *Os*phyC holoprotein is predominantly found as an *Os*phyB/phyC heterodimer *in vivo* (Xie *et al*., 2014), similar to that in *Arabidopsis* (Sharrock and Clack, 2004). *Os*phyC was not detected in the *OsphyB* mutant, indicating that phyB is essential for the expression and function of phyC in rice (Xie *et al*., 2014), as in *Arabidopsis*. Interestingly, de-etiolation was observed for the *OsphyA phyB* double mutant under Rc and FRc when nonphotoactivatable *Os*PHYB was introduced, although the double mutant did not respond to R or FR (Xie *et al*., 2014). This result indicates that phyC responds to R and FR in the presence of the phyB apoprotein without a chromophore in rice. Taken together, these studies revealed that phyC performs distinct functions from those of phyA and phyB in monocot and dicot plants.

Furthermore, some studies have proposed that phyC exhibits different absorption properties from those of phyA and phyB. Previously, only phyA was extracted from etiolated oat, pea and corn seedlings since phyA is abundant in these tissues and is used for spectroscopic studies. The absorption spectra of phyA prepared from etiolated seedlings showed absorption maximum (λ_max_) of 665-668 nm (Pr) and 726-730 nm (Pfr) (Everett and Briggs, 1970; Vierstra and Quail, 1983; Yamamoto and Furuya, 1983). Later, Remberg *et al*. (1998) reconstituted the N-terminal half-apoproteins of potato phyA and phyB with PΦB, measured the absorption spectra and attained λ_max_ values of 664 (Pr) and 729 (Pfr) nm for phyA and 668 (Pr) and 726 (Pfr) nm for phyB, respectively. Then, *At*phyB PSM holo-polypeptides were obtained by coexpression with PΦB synthesis enzymes. A λ_max_ of 663 (Pr) and 725 (Pfr) nm was attained for the chromopeptide with residues 1-624 (Zhang *et al*., 2013), and λ_max_ values of 664 (Pr) and 725 (Pfr) nm were attained for residues 90-624 lacking an N-terminal fragment (Burgie *et al*., 2014). Another reconstitution study with PΦB reported that the λ_max_ of *At*phyA, *At*phyB, *At*phyC and *At*phyE holopeptides in the Pr were 670, 669, 661 and 670 nm, respectively (Eichenberg *et al*., 2000). Thus, a hypsochromic shift occurred in the spectra of phyC compared with the other phytochromes, although it was difficult to precisely determine the λ_max_ due to the high noise level in the absorption difference spectra. Similarly, λ_max_ values of 650 and 721 nm were attained for the Pr and Pfr, respectively, in absorption spectra of a crude extract from phyC-GFP-overexpressing rice seedlings that harbored phyA- and phyB-deficient mutants (Xie et al., 2014); thus, compared to phyA and phyB in rice, phyC in the Pr absorbs approximately 10 nm less light. Burgie *et al*. (2021) reported the absorption spectra of phytochrome PSMs of *Arabidopsis*, potato and maize using a PΦB coexpression system in *E. coli*. The λ_max_ values attained for the phyC PSM polypeptides of *Arabidopsis*, potato and maize were 654, 654 and 655 nm, respectively, which were hypsochromically shifted compared to those of other phytochrome species. Taken together, these results suggest that the λ_max_ of phyC in monocot and dicot plants underwent a hypsochromic shift in the Pr.

In the present study, we prepared a photoactive PSM with a monocot plant, *Zea mays* (*Zm*), and dicot plants, *Solanum lycopersicum* (*Sl*) and *Arabidopsis thaliana*, using a coexpression system with PΦB synthetic genes. Then, we confirmed that the λ_max_ of phyC in the Pr undergoes a hypsochromic shift in monocot and dicot plants. The present amino acid substitution studies revealed that the GAF domain is involved in the shift, but the amino acid responsible for the shift was not identified.

To determine how the shift influences physiological functions, we compared the inhibition of hypocotyl growth under different R/FR ratios between the WT and phyB- or phyC-deficient mutants and found that phyC plays a complementary role under low R/FR conditions (less than 2.0). Since the activation peak of the phyC PSM from Pr to Pfr photoconversion shifted to a shorter wavelength than that of phyB, phyC can produce more Pfr under light conditions, which may explain the complementation. Furthermore, the absorption spectrum of phyC in the Pr was compared with the natural and leaf-filtered sunlight spectra, suggesting that complementation can effectively activate phytochromes under both light conditions in the field. This activity may be significant, especially in the higher latitude areas in which the R/FR decreases due to the longer passage through the atmosphere. This topic is discussed in relation to the presence of phyC variations that are dependent on latitudinal clines in the Northern Hemisphere.

## Materials and Methods

### Plasmid construction

A pTYB2-based expression vector for *At*phyB PSM polypeptides (Fig. 1A) was kindly provided by Dr. Matsushita at Kyoto University (Oka *et al*., 2004). *Arabidopsis* plants were grown at 22℃ under 16 h light/8 h dark conditions for 2 weeks, after which total RNA was purified from the leaf tissues to prepare a cDNA library. The PSM regions of the *AtphyA* and *AtphyC* genes corresponding to that of *AtphyB* were cloned from the cDNA library using the primer sets listed in Supplementary Table S1. cDNA libraries of the miniature tomato *Solanum lycopersicum* cultivar Micro-Tom and *Zea mays* var. *saccharata* were prepared from the leaf tissues following the same procedures used for *Arabidopsis*. The PSM regions of the *SlphyA*, *SlphyB2* and *SlphyF* genes (Hauser *et al*., 1995) and of the *ZmphyA2*, *ZmphyB1* and *ZmphyC1* genes (Sheehan *et al*., 2004), which correspond to those of *AtphyB*, were cloned from the cDNA libraries using the primer sets listed in Supplementary Table S1. The PSM regions of the *A. thaliana*, *S. lycopersicum* and *Z. mays* genes were inserted into the pTYB2 vector (New England Biolabs), which was previously digested with *Nde*I and *Sma*I for fusion with an Intein/CBD sequence, using the In-Fusion Cloning system (Takara-Bio).

Single amino acid substitutions of the *At*phy PSM polypeptides (Q233R/*At*phyB, A340S/*At*phyA, S335A/*At*phyC, and S621E/*At*phyB) and an *At*phyC hairpin substitution of the *At*phyB PSM polypeptide (hairpinC/*At*phyB) were generated using the specific primer sets listed in Table S1. To construct GAF-substituted polypeptides (C-GAF/*At*phyB and B-GAF/*At*phyC), pTYB2 vectors including the PSMs of *At*phyB and *At*phyC and the GAF regions of *At*phyB and *At*phyC were amplified, cloned and inserted into pTYB2 using the corresponding primer sets listed in Supplementary Table S1. Then, the overlapping sequences were ligated using the In-Fusion system and cloned.

### Preparation of PSM polypeptides

A pTYB2-based vector for the expression of PSM polypeptides and a pCDF-based vector that contained *heme oxygenase 1* and *phytochromobilin synthase* genes for the synthesis of PΦB (Muramoto *et al*., 1999; Kohchi *et al*., 2001) were transformed into the *Escherichia coli* Rosetta 2 strain (Merck). *E. coli* cells expressing PSM polypeptides and heme oxygenase 1 and phytochromobilin synthase were grown in LB supplemented with 20 μg mL^-1^ ampicillin, 20 μg mL^-1^ spectinomycin, and 10 μg mL^-1^ chloramphenicol at 37℃ until the OD_600 nm_ reached 0.2-0.3. Then, ampicillin and isopropyl-β-D-thiogalactopyranoside were added to final concentrations of 40 μg mL^-1^ and 1 mM, respectively. The culture was continued at 18℃ for 18 h in the dark. The following purifications were carried out under dim green light. Harvested *E. coli* cells were lysed in HEPES buffer (10 mM HEPES-NaOH, 500 mM NaCl, and 1 mM Na_2_EDTA, pH 7.8) containing 1 mM phenylmethylsulfonyl fluoride, and the supernatant was mixed and incubated with chitin beads (New England Biolabs). The PSM polypeptides were eluted with 50 mM DTT to cleave the intein. The eluate was applied to a Sephacryl S-100 HR column (GE Healthcare), equilibrated and eluted with the HEPES buffer. Elution profiles were monitored by the band patterns on SDS-PAGE gels, which were visualized by Coomassie brilliant blue staining and zinc fluorescence (Berkelman and Lagarias, 1986). The fractions containing the PSM polypeptides were collected, concentrated by ultrafiltration with Amicon Ultra-15 50,000 NMWL (Merck) (Figure S1) and used for spectroscopic analysis.

### Spectroscopic measurements

UV-visible absorption spectra of the PSM polypeptide solutions were measured at 25℃ with a spectrophotometer (3310, Hitachi-hitech) equipped with a thermoelectric cell holder (131-0305, Hitachi-hitech). For actinic irradiation of the sample solution, either an R light-emitting diode (LED) with λ_max_ = 657 nm (ISL-305X302-RRRR, CCS) or an FR LED with λ_max_ = 728 nm, (ISL-305X302-FFFF, CCS) was used at 30 μmol m^-2^ s^-1^ for 6 min. The intensity of the excitation light was measured with a photometric sensor (LI-210R, LI-COR), and all the procedures were performed in dim green light.

Action spectra of Pfr formation by R irradiation were measured using the excitation beam of a fluorescence spectrophotometer (RF5300, Shimadzu) as an actinic light source. An excitation beam with a 5 nm bandwidth was guided to the sample in the spectrophotometer through a quartz light guide (Φ = 1 cm × 1 m). After irradiation of PSM samples with FR (peak at 720 nm, 15 nm band width) for 5 minutes to convert PSM samples into Pr and absorption spectra were recorded. Then, PSM samples were irradiated with R every 10 nm in the 620 to 650 nm and 670 to 690 nm region and every 5 nm in the 650 to 670 nm region for 2 minutes to saturate Pfr formation. Using absorption spectra of PSM samples before R irradiation and their absorption difference spectra between before and after R irradiation at each activation wavelength, the Pfr formation efficiency was calculated as the ratio of the absolute value of λ_max_ absorption of Pfr produced by R irradiation to the value of λ_max_ absorption of Pr before R irradiation. The values obtained from the Pfr formation efficiencies fit a regression curve (R^2^=0.95 for *Sl*phyB2 and *Sl*phyF).

Solar spectra and transmission spectra of sunlight passing through one or two leaves of *Arabidopsis* plants were measured in Sakai, Osaka, Japan (lat. 34° 54’ N. and long. 135° 50’ E); these tests were performed in April at noon on a clear day using a spectroradiometer HR2000+ (Ocean Optics).

### Clustering analysis

Amino acid sequences of full-length phytochromes were obtained from the NCBI database. The corresponding PSM sequences were compared with those of *At*phyB. Multiple sequence alignments were performed with ClustalX.

### Plant materials and hypocotyl growth conditions

*Arabidopsis thaliana* Columbia-0 (Col-0) and its mutants (*phyB-9* and *phyC-2*) in the Col-0 background (Monte *et al*., 2003; Effendi *et al*., 2013) were used in this study. For hypocotyl analysis, the seeds were sterilized using 70% ethanol, sown on 0.3% Gelrite plates supplemented with 1× Murashige and Skoog media without sucrose and stratified at 4℃ in the dark for 3 days. Seeds were grown under continuous irradiation with a white light LED (LDA5L-G AG52, OHM electric Inc.), which emits 0.074 μmol m^-2^ s^-1^ at 660 nm and 0.010 μmol m^-2^ s^-1^ at 734 nm light, as well as a FR with an emission maximum at 734 nm (3 in 1 LED, NK system). The output power of the FR LEDs was 100, 75, 50, 25 and 0% to achieve R/FR ratios of 0.5, 0.7, 1.0, 2.0 and 10, respectively (Supplementary Fig. S2). After 4 days of incubation, hypocotyl lengths were measured using ImageJ. The average hypocotyl lengths were calculated from 3 sets of tests, and data from 3 groups with more than 50 seedlings were used for each test. The fluorescence rates were measured with a spectrophotometer (S-2440 model II, NK system).

## Results

### The hypsochromically shifted λ_max_ of phyC in the Pr is conserved in dicot and monocot plants compared with those of phyA and phyB

To confirm whether the hypsochromic shift observed for phyC in the Pr is conserved among angiosperms, holo-PSM polypeptides of phytochromes corresponding to an *At*phyB (residues 1-651) were prepared from *Arabidopsis thaliana* (*At*phys), *Solanum lycopersicum* (*Sl*phys), and *Zea mays* (*Zm*phys) as photoactive PΦB adducts (Supplementary Fig. S1).

The λ_max_ values of *At*phyA and *At*phyB PSMs were found at 662 and 663 nm in the Pr and at 727 and 724 nm in the Pfr, respectively (Fig. 1B; Table 1). In contrast, the λ_max_ value obtained for the Pr of *At*phyC PSM was 654 nm, which is 8 and 9 nm shorter than those of the *At*phyA and *At*phyB PSMs, respectively, while the λ_max_ of Pfr was 725 nm, which was not significantly different among the three PSMs (Fig. 1B; Table 1). PSM polypeptides were also prepared from a dicot tomato and a monocot maize. *S. lycopersicum* has five phytochrome genes, *phyA*, *phyB1*, *phyB2*, *phyE*, and *phyF* (Hauser *et al*., 1995). A phylogenetic analysis of full-length amino acid sequences classified *Sl*phyF into a phyC group (Alba *et al*., 2000). Since both *Sl*phyB1 and *Sl*phyB2 function in fruit development (Gupta *et al*., 2014), *Sl*phyB2 was used for the present study. The PSM of the tomato phytochromes displayed reversible photoconversion by R and FR irradiation (Fig. 1B). The absorption difference spectra of the *Sl*phyA and *Sl*phyB2 PSMs showed λ_max_ in the Pr at 662 and 668 nm, respectively. The λ_max_ of *Sl*phyC was 657 nm, showing 5 and 11 nm hypsochromic shifts compared with those of *Sl*phyA and *Sl*phyB2, respectively. In contrast, the λ_max_ values in the Pfr of *Sl*phyA, *Sl*phyB2, and *Sl*phyF were 723, 724, and 721 nm, respectively, indicating no significant differences (Table 1).

**Table 1.** Absorption maxima of the phytochrome PSMs in the Pr and Pfr.

| | Pr $\lambda_{\text{max}}$ [nm] | Pfr $\lambda_{\text{max}}$ [nm] |
| --- | --- | --- |
| *At*phyA | 662 | 727 |
| *At*phyB | 663 | 724 |
| *At*phyC | 654 | 725 |
| *Sl*phyA | 662 | 723 |
| *Sl*phyB2 | 668 | 724 |
| *Sl*phyF | 657 | 721 |
| *Zm*phyA2 | 664 | 724 |
| *Zm*phyB1 | 662 | 724 |
| *Zm*phyC1 | 654 | 720 |
| Q233R/*At*phyB | 659 | 716 |
| A340S/*At*phyA | 661 | 722 |
| S335A/*At*phyC | 654 | 726 |
| S621E/*At*phyB | 656 | 717 |
| hairpinC/*At*phyB | 661 | 714 |
| C-GAF/*At*phyB | 652 | 722 |
| B-GAF/*At*phyC | 660 | 722 |

Table 1 Absorption maxima of the phytochrome PSMs in the Pr and Pfr.
| | Pr $\lambda_{\text{max}}$ [nm] | Pfr $\lambda_{\text{max}}$ [nm] |
| --- | --- | --- |
| *AtphyA* | 662 | 727 |
| *AtphyB* | 663 | 724 |
| *AtphyC* | 654 | 725 |
| *SlphyA* | 662 | 723 |
| *SlphyB2* | 668 | 724 |
| *SlphyF* | 657 | 721 |
| *ZmphyA2* | 664 | 724 |
| *ZmphyB1* | 662 | 724 |
| *ZmphyC1* | 654 | 720 |
| Q233R/*AtphyB* | 659 | 716 |
| A340S/*AtphyA* | 661 | 722 |
| S335A/*AtphyC* | 654 | 726 |
| S621E/*AtphyB* | 656 | 717 |
| hairpinC/*AtphyB* | 661 | 714 |
| C-GAF/*AtphyB* | 652 | 722 |
| B-GAF/*AtphyC* | 660 | 722 |

The tetraploid maize genome contains duplicates of phyA, phyB, and phyC, and all six genes were transcribed in the seedlings, in which phyA1, phyB1, and phyC1 were predominantly expressed in planta (Sheehan *et al*., 2004). In the present study, the PSMs of *Zm*phyA2, *Zm*phyB1, and *Zm*phyC1 were used. All the PSMs of *Zm*phyA2, *Zm*phyB1 and *Zm*phyC1 exhibited changes in absorption between the Pr and the Pfr upon R and FR irradiation (Fig. 1B) and showed a λ_max_ in the Pr at 664, 662, and 654 nm, respectively; therefore, a hypsochromic shift occurred for *Zm*phyC1, while their λ_max_ in the Pfr were at 724, 724, and 720 nm, respectively (Table 1).

The λ_max_ values listed in Table 1 indicate that phyC has a hypsochromically shifted Pr λ_max_ compared with phyA and phyB, while their Pfr λ_max_ values are comparable in monocot and dicot plants. This finding suggests that the absorption traits of phyC are ubiquitous among angiosperms.

### phyC-specific single amino acid residue substitution does not reproduce the hypsochromically shifted λ_max_ of phyC in Pr

First, we compared the amino acid sequences of PSMs among the phyA, phyB, and phyC groups to determine the amino acid residues responsible for phyC absorption. The following phyC-specific amino acid residues in the PSM were detected: R194, S335 and E580 in *At*phyC (Supplementary Fig. S4, letters with a green background). However, based on the crystal structure of an NTS-lacking *At*phyB PSM (residues 90-624, PDB: 4OUR, Burgie *et al*., 2014), these three amino acid residues are not located within the direct interaction distance of PΦB (Fig. S5). These residues were substituted between *At*phyC and *At*phyA or *At*phyB.

R194 is conserved in basic Arg or Lys among phyC groups, while Gln is conserved among phyA- and phyB-type phytochromes (Supplementary Fig. S4). The λ_max_ of Q233R/*At*phyB, in which Q233 of *At*phyB corresponding to R194 of *At*phyC is replaced by Arg, was 659 nm for Pr and 716 nm for Pfr (Table 1; Fig. 2), both of which were shifted to slightly shorter wavelengths than those of *At*phyB. S335 *At*phyC is present in near and downstream of the chromophore-binding Cys residue, and Ala is highly conserved among phyA- and phyB-type phytochromes (Supplementary Fig. S4). The λ_max_ of A340S/*At*phyA, in which A340 in *At*phyA (corresponding to S335 in *At*phyC) is replaced by Ser, are 661 nm for Pr and 722 nm for Pfr (Table 1; Fig. 2). The Pfr of A340S/*At*phyA shifted the λ_max_ to a 5 nm shorter wavelength; however, that of Pr was hardly affected. Similarly, λ_max_ of *At*phyC was not significantly different for S335A/*At*phyC, in which S335 of *At*phyC was replaced by Ala (Table 1; Fig. 2). The λ_max_ of S621E/*At*phyB, in which S621 of *At*phyB corresponding to E580 of *At*phyC is replaced by Glu, is 656 nm for Pr and 717 nm for Pfr (Table 1; Fig. 2). The S621E substitution shifted the λ_max_ of *At*phyB in Pr to a shorter wavelength, similar to *At*phyC; however, the substitution also shifted the λ_max_ of Pfr, which did not precisely match to the absorption property of *At*phyC.

**Fig. 2.**
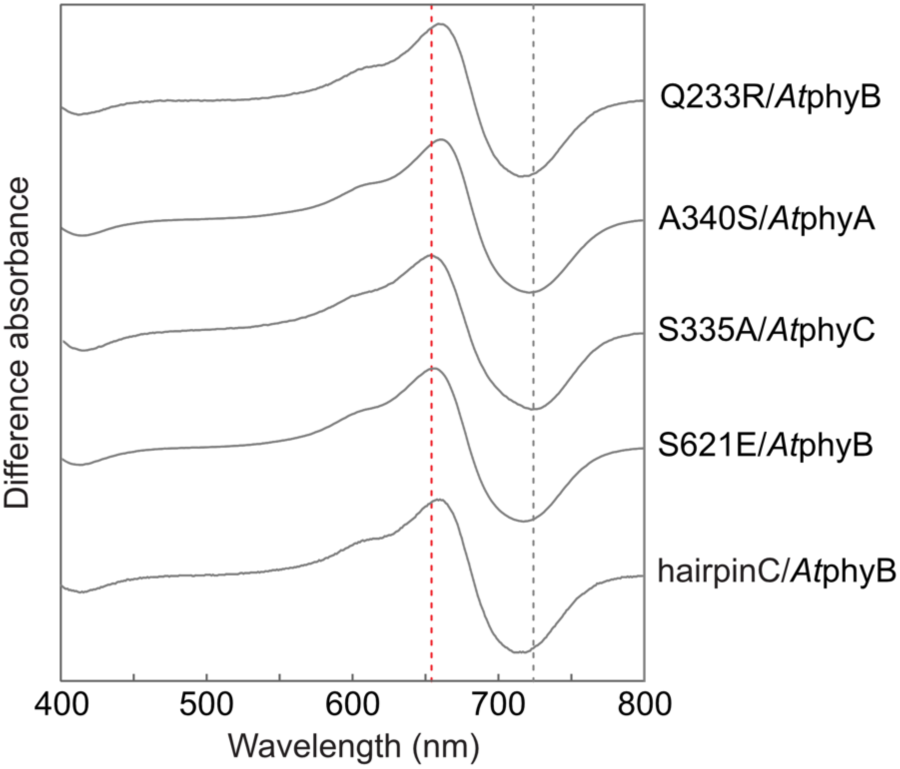
Absorption difference spectra of amino acid-substituted PSMs of *Arabidopsis* phytochromes between Pr and Pfr. The names of the spectra indicate locations of mutations studied in the present study. The hairpinC indicates the phyC-specific hairpin region substituted in this study. The red and gray dashed lines indicate the position of the λ_max_ of *At*phyC in the Pr and Pfr, respectively.

### The hairpin does not contribute to the hypsochromically shifted λ_max_ of phyC in the Pr

Crystal structures of plant, cyanobacteria and proteobacteria phytochromes show that the surface of the chromophore-binding pocket in the GAF domain is covered with a loop, which is a part of the structure called a “tongue” or “hairpin” that extends from a core of the PHY domain (Essen *et al*., 2008; Burgie *et al*., 2014) (Fig. 1A; Supplementary Fig. S4, Fig. S5). Substituting highly conserved amino acid residues in the *At*phyB hairpin led to hypsochromic shifts in the λ_max_ in Pr, while the λ_max_ values in Pfr were unaffected (G564-E, WGG-SEE, R582-A in Burgie *et al*., 2014) (Supplementary Table S2). To determine whether the residues in the hairpin are involved in phyC-specific absorption, ten amino acid residues of *At*phyC from R527 to K536 (Supplementary Fig. S4, letters with violet background) were introduced to the corresponding region of *At*phyB (hairpinC/*At*phyB) (Table 1; Fig. 2). The hairpinC/*At*phyB PSM exhibited photoconversion between the Pr and Pfr (data not shown). The λ_max_ in Pr was 661 nm and close to that of the *At*phyB (663 nm), while the λ_max_ in Pfr shifted from 724 to 714 nm, similar to that of a S621E/*At*phyB substitute (Table 1; Fig. 2). Accordingly, the amino acids in this hairpin loop region may not be involved in maintaining the phyC-specific absorption property.

### Hypsochromically shifted λ_max_ of phyC in Pr is dependent on the GAF domain

The GAF domain is located at the center of the PSM in a phytochrome molecule and binds a chromophore PΦB at a conserved Cys residue (Fig. 1A; Supplementary Fig. S4). To investigate the role of the GAF domain in maintaining phyC-specific absorption properties, the GAF domains were interchanged between the PSMs of *At*phyB and *At*phyC. “C-GAF/*At*phyB” and “B-GAF/*At*phyC” represent “the PSM of *At*phyB with *At*phyC-GAF” and “the PSM of *At*phyC with *At*phyB-GAF”, respectively. C-GAF/*At*phyB and B-GAF/*At*phyC showed photoconversion from Pr to Pfr (Fig. 3A, B). Introducing phyC-GAF to phyB converted the λ_max_ of phyB in Pr (663 nm) to a phyC-like peak (652 nm) and vice versa (654 to 660 nm), while it slightly altered the λ_max_ of Pfr (C-GAF/*At*phyB; from 724 to 722 nm, B-GAF/*At*phyC; from 725 to 722 nm) (Table 1; Fig. 3A-C). Thus, the recipient phytochrome between phyB and phyC can obtain the native λ_max_ of *At*phy-GAF, while the Pfr λ_max_ remains almost unchanged, indicating that the GAF domain plays a significant role in maintaining the phyC-specific absorption trait.

**Fig. 3.**
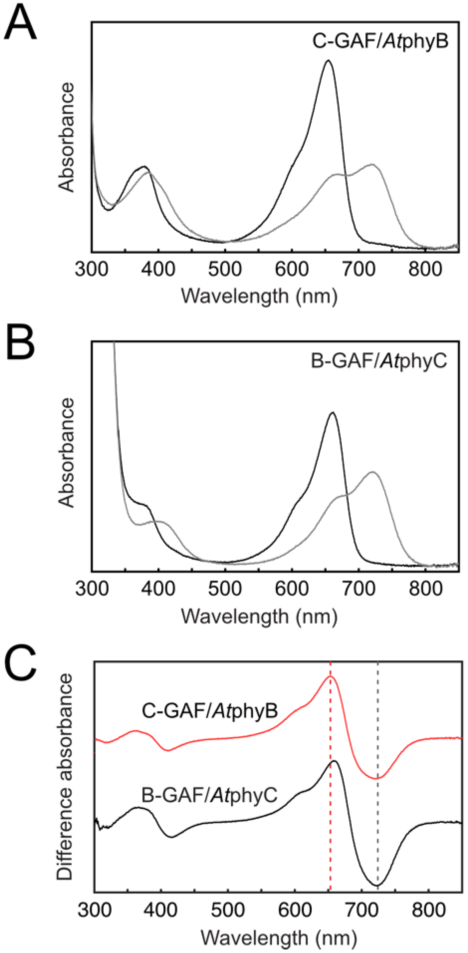
Pr and Pfr absorption spectra of the GAF domain interchanged with phyB and phyC. (A) Pr and Pfr absorption spectra of C-GAF/*At*phyB. C-GAF/*At*phyB represents the PSM of *At*phyB with *At*phyC GAF. (B) Pr and Pfr absorption spectra of B-GAF/*At*phyC. (C) Their absorption difference spectra between Pr and Pfr. The red and gray dashed lines indicate the position of the λ_max_ of *At*phyC in the Pr and Pfr, respectively.

### Activation spectra of tomato phyB2 and phyF PSMs for the photoconversion from Pr to Pfr

Based on the hypsochromically shifted λ_max_ of the phyC PSM in Pr, the Pr of phyC undergoes maximum photoconversion to Pfr at a shorter wavelength than that of phyA and phyB. To examine this possibility, the activation spectra for the photoconversion of *Sl*phyB2 and *Sl*phyF from Pr to Pfr in the R region from 620 to 690 nm were measured and compared (Fig. 4). *Sl*phyB2 showed maximum photoconversion at 665 nm, while *Sl*phyF showed maximum photoconversion at 650 nm, consistent with the λ_max_ of *Sl*phyB2 in Pr (668 nm) and that of *Sl*phyF in Pr (657 nm) (Table 1; Fig. 1B). Given that the quantum yield of the photoconversion from Pr to Pfr of PSM is not very different between the two phytochromes (Burgie *et al*., 2021), the hypsochromically shifted Pr absorption peak of phyC may enable a plant to sense R at a shorter wavelength than the other phyB and phyA *in vivo*.

**Fig. 4.**
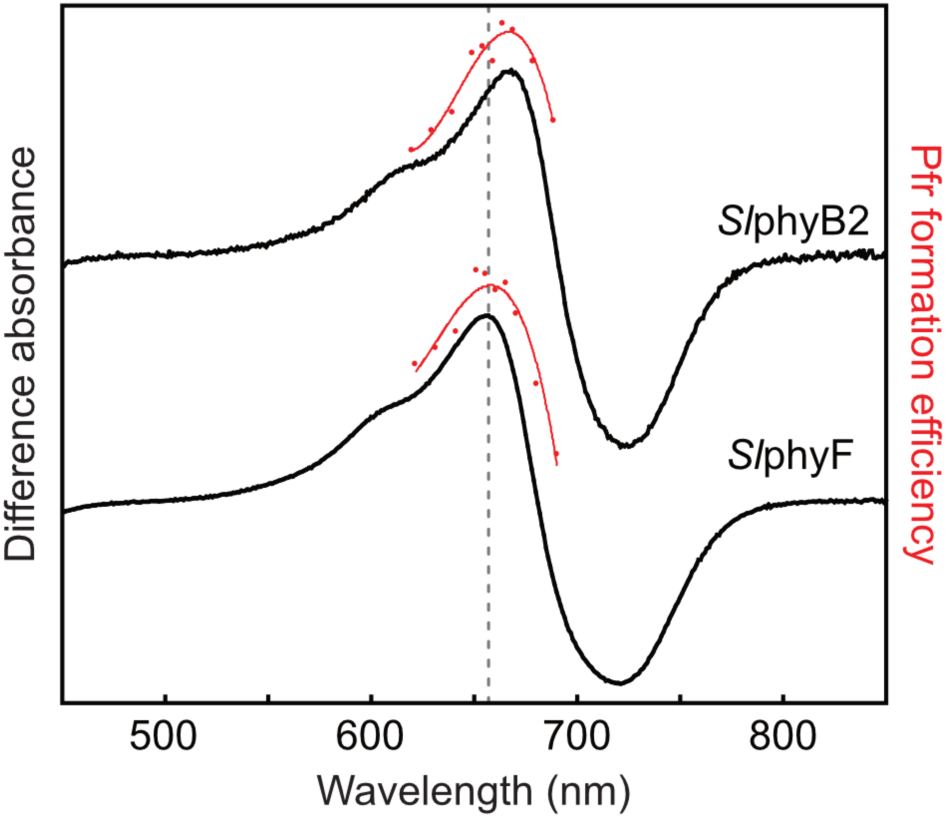
Activation spectra of *Sl*phyB2 and *Sl*phyF for the photoconversion of Pr to Pfr. The red dots and the red lines indicate Pfr formation efficiencies at each wavelength and regression curves (R^2^=0.95 for *Sl*phyB2 and *Sl*phyF), respectively. The black lines show their absorption difference spectra between Pr and Pfr, which are the same as the spectra shown in Fig. 1B. The black dashed line indicates the position of the Pr λ_max_ of *Sl*phyF.

### Contribution of phyC to the inhibition of hypocotyl elongation at lower R/FR

To determine the physiological implications of the hypsochromic shift in the Pr absorption of phyC, the inhibition of hypocotyl elongation was compared among the WT, *PHYB* mutant and *PHYC* mutant of *Arabidopsis* under various R (660 nm) /FR (734 nm) wavelengths. The ratio was controlled to 0.5, 0.7, 1.0, 2,0 or 10 by adding the emission light from the FR LED to the white light LED (Supplementary Fig. S2). Col-0, *phyB-9* and *phyC-2* mutants in the Col-0 background were used as WT, *phyB* and *phyC* mutants since the *PHYC* gene of *A. thaliana* ecotype L*er* contains at least ten nonsynonymous changes, suggesting the limited activity of L*er* phyC.

In the WT, hypocotyl elongation was inhibited to a similar extent at all the R/FR ratios, except that the inhibition was slightly lower at R/FR 0.5 (Fig. 5 Col-0). The inhibition of *phyB-9* was eliminated; however, the reduction in inhibition was slightly less at an R/FR of 0.5 compared to that at an R/FR of 0.7 -10 (Fig. 5 *phyB-9*). Interestingly, the inhibition of *phyC-2* was enhanced at an R/FR of 0.5-2.0 compared to that of Col-0 and *phyB-9* (Fig. 5), indicating that phyC contributed to the inhibition. This contribution of phyC decreased as the R/FR increased from 0.5 to 2.0. At an R/FR of 2.0, the inhibition reached almost the same level as that of Col-0, possibly due to the prevalent contribution of phyB at this R/FR. These data indicate that phyC plays a complementary role in inhibiting hypocotyl elongation at low R/FR ratios, at which point phyB functions insufficiently.

**Fig. 5.**
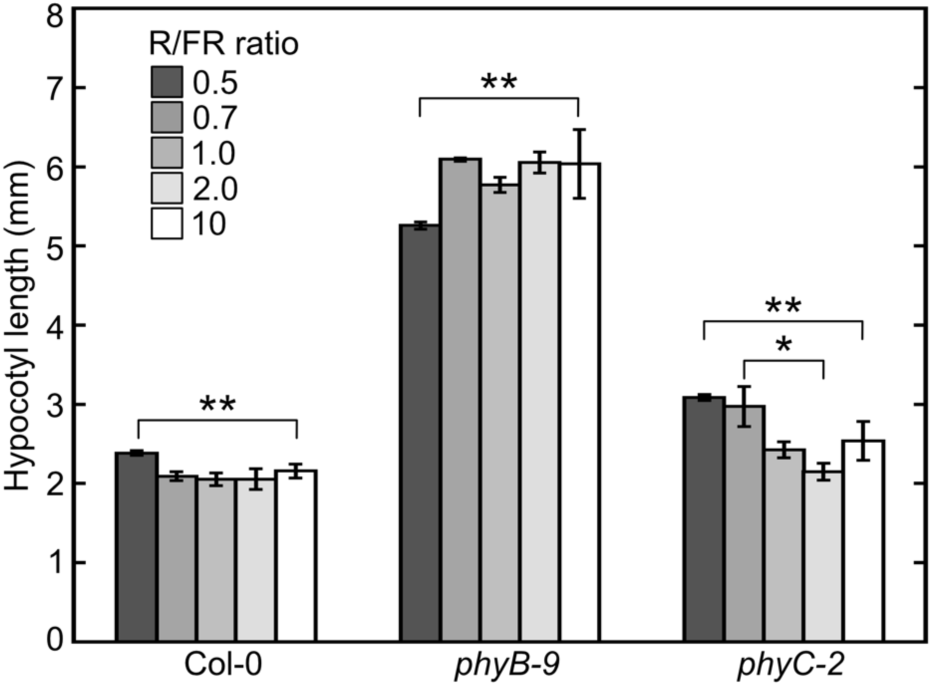
Hypocotyl length of *Arabidopsis* seedlings of a wild type, Col-0, and *phyB-9* and *phyC-2* mutants, after 4 days of growth under different R/FR ratios from 0.5 to 10. The bars represent the standard errors of the values. The average hypocotyl lengths were calculated from 3 sets of tests, and data from 3 groups with more than 50 seedlings were used for each test. One and two asterisks represent *p* < 0.05 and *p* < 0.01, respectively, according to Student’s t test.

## Discussion

### The hypsochromically shifted λ_max_ of phyC in the Pr is conserved in monocot and dicot plants

In this study, we revealed that the λ_max_ values of PSM of phyC from *Arabidopsis thaliana*, *Solanum lycopersicum* and *Zea mays* were hypsochromically shifted in the Pr by approximately 10 nm compared with those of phyA and phyB; in contrast, the λ_max_ values in the Pfr were not significantly different among the three plants (Table 1; Fig. 1B). Compared with the other *At*phys, the PΦB-reconstituted PSM of *At*phyC has a hypsochromically shifted Pr λ_max_ (Eichenberg *et al*., 2000). The addition of a regulatory module to a PSM does not alter the λ_max_ of *At*phyB in Pr or Pfr, although it modifies the reaction rates of thermal reversion from Pfr to Pr (Burgie *et al*., 2017); therefore, the present hypsochromic shifts observed in the PSM may be preserved in their full-length phytochromes. In addition to the present and reported results obtained by *in vitro* systems, the absorption difference spectra of the crude extract from *phyA phyB*-deficient rice overexpressing phyC-GFP revealed that the λ_max_ was hypsochromically shifted in Pr (Xie *et al*., 2014). These results indicate that the phyC-specific hypsochromic shift in Pr λ_max_ is preserved in monocot and dicot plants and may be conserved among angiosperms.

### Amino acid residues responsible for the hypsochromically shifted λ_max_ of phyC in the Pr

The present study revealed three phyC-specific amino acid residues, R194, S335, and E580 (letters with a green background in Supplementary Fig. S4; Supplementary Fig. S5); however, none of the single amino acid substitutions between *At*phyC and *At*phyA or between *At*phyC and *At*phyB caused the hypsochromically shifted λ_max_ of phyC to phyA or phyB and vice versa (Table 1; Fig. 2). No phyC-specific residue is included in the ten amino acid residues that interact with PΦB through hydrogen bonding and van der Waals interactions in the crystal structure of the truncated *At*phyB PSM (Burgie *et al*., 2014) (Supplementary Fig. S4, letters with a cyan background; Supplementary Fig. S5, residues shown with sticks in cyan). Amino acid substitution studies of this truncated *At*phyB PSM revealed that substitution at six amino acid residues (Y104 and I108 in the PAS domain, R352 and S370 in the GAF domain, and G564 and R582 in the PHY domain) caused the λ_max_ to hypsochromically shift in the Pr and led to a normal photoconversion to Pfr, although small hypsochromic shifts were observed for the Pfr λ_max_ (Oka et al., 2008; Zhang *et al*., 2013; Burgie *et al*., 2014) (red arrows in Supplementary Fig. S4; Table S2).

The phyC-specific R194 corresponding to Q233 of *At*phyB resides on a helix between the PAS and the GAF domains (Supplementary Fig. S4; Fig. S5), and no amino acid residue with red arrows is found nearby (Supplementary Fig. S4). Q233 is located at the interface between the two subunits of a dimeric phyB crystal structure (Burgie *et al*., 2014) and is far from the PΦB chromophore (Supplementary Fig. S5). This may explain the inability of the amino acid substitution, which changes the λ_max_ of PΦB. S335, corresponding to *At*phyB A374, is located in the middle of the β5 strand, which is a component of the β-sheets behind the PΦB chromophore and is next to the α5 helix that binds PΦB in the GAF domain (Supplementary Fig. S4; Fig. S5). S370 is near the A374, and substituting S370 to Phe is reported to induce a hypsochromic shift of λ_max_ in Pr and a normal photoconversion to Pfr (Oka *et al*., 2008) (Supplementary Table S2; a pink background and red arrow in Fig. S4). S370 is close to the C-ring propionate of PΦB, which may play a role in the conformational change in phyB from Pr to Pfr (Burgie *et al*., 2014). Therefore, S335 of phyC at a distance from PΦB could contribute to the hypsochromic shift; however, the single amino acid substitution of *At*phyB did not result in this shift. E580 of *At*phyC, which corresponds to S621 of *At*phyB, was not resolved in the crystal structure, as described in the Results. Substitution of *At*phyB S621 with Glu hypsochromically shifted the λ_max_ of Pr and Pfr (Table 1; Fig. 2). Based on the crystal structure and cryoelectron microscopy structure (Li *et al*., 2022), S621 may be far from PΦB in a phyB molecule. Interestingly, this distant amino acid can affect the λ_max_ of PΦB. The region that includes S621 may be involved in some conformational changes that accompany phototransformation from Pr to Pfr, which modulates the coplanarity of the π-electron conjugating system of PΦB (Hanzawa *et al*., 2002). The phyC-specific residues found in the present study are located away from PΦB and may not play a critical role or not sufficiently work to hypsochromically shift the λ_max_ in the Pr and prevent the λ_max_ in the Pfr from changing.

### Contribution of the hairpin to the hypsochromically shifted λ_max_ of phyC in the Pr

The hairpin structure of *At*phyB resides between a β5 strand and a β6 strand in the C-terminal end region of the PHY domain and consists of two β (β_exit_ and β_int_) strands and a loop between them that interacts with the chromophore in the GAF domain (Supplementary Fig. S5). Replacing the center region of the phyB hairpin with that of phyC did not provide phyB with the absorption trait of phyC (Table 1; Fig. 2). The rotation and helical conversion of the strand-β_exit_ may induce positional exchange of the two strands and play important roles in signal transduction (Burgie *et al*., 2014), but may be the least important region for maintaining the absorption characteristics of phyC.

### Contribution of the GAF domain to maintain the hypsochromically shifted λ_max_ of phyC in the Pr

Exchange of the GAF domain between *At*phyB and *At*phyC instead of substituting amino acids led to absorption characteristics specific to each GAF domain, although the λ_max_ of Pfr showed a slight short-wavelength shift (Table 1; Fig. 3). The PΦB bound to B-GAF/*At*phyC was partially lost due to heat treatment during SDS-PAGE, but photoconversion between Pr and Pfr was successfully demonstrated (Fig. 3B, C; Supplementary Fig. S1). C-GAF/*At*phyB and B-GAF/*At*phyC mimicked their original GAF-derived Pr absorption characteristics. Plant, cyanobacterial and nonphotosynthetic eubacterial phytochromes require a PAS-GAF-PHY domain set for the photoreversible transition between Pr and Pfr and an NTE for the Pfr stabilization (Oka *et al*., 2004; Wagner *et al*., 2005; Essen *et al*., 2008; Burgie *et al*., 2014). The present result correspond with the previous belief that the GAF domain plays a major role in controlling the absorption spectral properties of phytochromes (Burgie and Vierstra, 2014).

### Involvement of phyC in the inhibition of hypocotyl elongation under low R/FR conditions

The present results showed that phyB plays a major role in the inhibition of hypocotyl elongation by R (Fig. 5, Col-0 and *phyB-9*), in accordance with previously reported results (Reed *et al*., 1994; Sánchez-Lamas *et al*., 2016). In addition to the role of phyB, phyC lead to a small inhibition under low R/FR conditions from 0.5 to 1.0 (Fig. 5, *phyC-2*). PhyC is known to show R-dependent inhibition of hypocotyl elongation (Flanklin *et al*. 2003; Monte *et al*. 2003). The present results revealed that phyC plays a complementary role to phyB in the inhibition of hypocotyl elongation under these low R/FR conditions. Considering the hypsochromically shifted maximum of the activation spectra from Pr to Pfr (Fig. 4), R can produce more Pfr in phyC than in phyB. Accordingly, the action of phyC Pfr may become more prominent than that of phyB under low R/FR conditions. Thus, the hypsochromically shifted λ_max_ could lead to the complementary role of phyC in the hypocotyl growth inhibition.

Another possible factor involved in the complementary role of phyC is thermal reversion from Pfr to Pr. The lifetime of Pfr is important for physiological function and is defined by the thermal reversion, which differs among phytochromes. The major component of the thermal reversion is reported to be 21 times faster in phyB PSM than in phyC PSM at 25℃ (Burgie *et al*., 2021). Coimmunoprecipitation analyses revealed that phyC functions as a phyB/phyC heterodimer *in vivo* (Sharrock and Clack, 2004), while phyB forms a phyB/phyB homodimer (Clack *et al*., 2009). Binding of phyC to phyB possibly delays the thermal reversion of the phyB/phyC heterodimer compared with the phyB/phyB homodimer, although no data are available. This slowed thermal reversion could be involved in the complementary role of phyC.

### Physiological implication of the hypsochromically shifted λ_max_ of phyC in the Pr

In the field, the hypsochromically shifted λ_max_ of phyC may be advantageous because more R can be obtained than phyB. The overlapping areas between the sunlight spectrum and the Pr absorption spectrum were 0.67 +/- 0.002 (SE) and 0.69 +/- 0.002 (SE) (arbitrary unit) for phyB and phyC, respectively, showing that phyC can absorb 4% more R in the field (Fig. S3). The intensity of R in sunlight is significantly reduced after passing through a leaf due to the strong R absorption of photosynthetic pigments (Franklin, 2008). According to our measurements, the R/FR ratio under sunlight decreased from 1.23 to 0.08 after passing through one leaf of *Arabidopsis* (Supplementary Fig. S3). The overlapping areas between the spectrum of sunlight transmitted through one leaf and the absorption spectrum of phyC is approximately 5% larger than that of phyB, suggesting that phyC produces 5% more active Pfr than phyB after passing through a leaf in the field. Within a single leaf, the quality of light reaching the cells may differ depending on the distance from the leaf surface. Compared with cells far from the leaf surface, cells near the leaf surface receive light with a greater R/FR ratio. In the latter case, 5% more activated phyC may assist the function of phyB. Thus, the hypsochromically shifted λ_max_ of phyC in the Pr may act as an effective tool to capture sunlight signals.

The R/FR ratio in the natural light reaching the ground decreases as the latitude increases due to the increased light path through the atmosphere, during which light with shorter wavelengths becomes more scattered (Wilbert *et al*., 2016). Accordingly, the R/FR is smaller in the northern high-latitude area than in the southern low-latitude area in the Northern Hemisphere. Interestingly, *PHYC* allele comparison of 163 *Arabidopsis* WT strains revealed a significant association between the locus and latitude in the Northern Hemisphere (Balasubramanian *et al*., 2006). *PHYC* alleles harboring mutations, such as a nonsense change to convert an amino acid-encoding codon to a stop codon (Fr-2 *PHYC* allele) or multiple mutations resulting in nonsynonymous changes (Fr-2 and L*er PHYC* alleles), tend to be distributed at southern latitudes. On the other hand, *Arabidopsis* WT strains with a functional *PHYC* allele (Col-0 *PHYC* allele) tend to appear at northern latitudes. *Arabidopsis* may develop the *PHYC* alleles to use the natural light that contains a lower R/FR ratio via a hypsochromic shift in the Pr. It is interesting to study the absorption spectral properties of these *PHYC* alleles to determine the correlation of the hypsochromic shift in the Pr with latitude. In addition to the latitudinal cline distribution of *PHYC* alleles, the contribution of phyC to short day flowering and the accompanying functional *FRIGIDA* (*FRI*) are also latitude dependent (Balasubramanian *et al*., 2006). Few natural alleles, such as those of *PHYC*, are involved in the phenotypic variation in both seedling growth and flowering time. (Aukerman *et al*., 1997; El-Assal *et al*., 2001; Maloof *et al*., 2001; Werner *et al*., 2005). Studying the role of phyC in the latitudinal cline of flowering time is also interesting.

## Supplementary data

The following supplementary data are available at JXB online.

Table S1. Primer sequences used for cloning phytochrome photosensory modules (PSMs) and making mutant constructs.

Table S2. Amino acid substitutions which induce hypsochromic shift of *At*phyB PSM in Pr.

Fig. S1. Images of SDS-PAGE and zinc-fluorescence of holo-PSM polypeptides.

Fig. S2 Emission spectra of LEDs used for inhibition of hypocotyl elongation analysis.

Fig. S3 Absorption spectra of phytochrome B and C in the Pr and emission spectra of sunlight.

Fig. S4 Amino acid sequence alignments of the PSMs of phytochromes.

Fig. S5 Three-dimensional structures of the *At*phyB PSMs with 90-624 residues (PDB: 4OUR).

## Acknowledgements

We would like to thank Dr. Matsushita at Kyoto University for giving pTYB2-based expression vector for *At*phyB PSM polypeptides (phyB_N651).

## Author contributions

SY conceptualized and performed absorption, plant phenotypic analyses. KO assisted with absorption analysis. SY and ST prepared the manuscript.

## Data availability

The data underlying this article are available in the article and in its online supplementary material.

## Abbreviations

At: Arabidopsis thaliana
FR: far-red light
FRc: continuous far-red light
GAF: cGMP phosphodiesterase/adenylyl cyclase/FhlA
HKRD: histidine kinase-related domain
LED: light-emitting diode
NTE: N-terminal extension Os Oryza sativa
PAS: Per-Arnt-Sim
Pfr: far-red light absorbing form
phy: phytochrome
Pr: red light absorbing form
PSM: photosensory module
PΦB: phytochromobilin
R: red light
Rc: continuous red light
Sl: Solanum lycopersicum
WT: wild type
Zm: Zea mays
λ_max_: absorption maximum

